# Using human sequencing to guide craniofacial research

**DOI:** 10.1101/356717

**Authors:** Ryan P. Liegel, Erin Finnerty, Lauren Ward, Andrew DiStasio, Robert B. Hufnagel, Howard M. Saal, Cynthia A. Prows, Rolf Stottmann

**Affiliations:** Division of Human Genetics; Division of Patient Services; Division of Developmental Biology; Department of Pediatrics, Cincinnati Children’s Hospital Medical Center University of Cincinnati College of Medicine, Cincinnati, OH Cincinnati, OH 45229; Shriners Hospitals for Children, Cincinnati

## Abstract

A recent convergence of technological innovations has re-energized the ability to apply genetics to research in human craniofacial development. Next-generation exome and whole genome sequencing have dropped in price significantly making it relatively trivial to sequence and analyze patients and families with congenital craniofacial anomalies. A concurrent revolution in genome editing with the use of the CRISPR-Cas9 system enables the rapid generation of animal models, including mouse, which can precisely recapitulate human variants. Here we summarize the choices currently available to the research community. We illustrate this approach with the study of a family with a novel craniofacial syndrome with dominant inheritance pattern. The genomic analysis suggest a causal variant in *AMOTL1* which we modeled in mice. We also made a novel deletion allele of *Amotl1*. Our results indicate that *Amotl1* is not required in the mouse for survival to weaning. Mice carrying the variant identified in the human sequencing studies, however, do not survive to weaning in normal ratios. The cause of death is not understood for these mice complicating our conclusions about the pathogenicity in the index patient. Thus, we highlight some of the powerful opportunities and confounding factors confronting current craniofacial genetic research.

## INTRODUCTION

During the first trimester of human development, multiple tissue prominences are created which then grow and fuse to form the early elements of the face. Proper formation of the craniofacial tissues requires coordination of multiple cell types of different embryonic origins to ultimately produce the morphologically complex mature craniofacial structures. A particularly critical cell type for craniofacial development is the neural crest cell (NCC) population. NCCs are born at the interface of the neural and surface ectoderm and migrate to multiple positions in the body and contribute to the majority of the craniofacial structures. Disruption of proper craniofacial development leads to craniofacial anomalies which are some of the most common congenital anomalies with a collective incidence of approximately 1/600 live births (‘Global strategies to reduce the health care burden of craniofacial anomalies: report of WHO meetings on international collaborative research on craniofacial anomalies,’ 2004). These anomalies are typically divided into two main classifications: isolated cleft lip and/or cleft palate or, alternatively, syndromic malformations where additional tissues are affected. A caveat to these classifications is, of course, that the undiscovered phenotypes in any specific patient may or may not be related to a specific variant causing the craniofacial anomaly originally called an isolated cleft, especially given all the lineages NCCs contribute to.

The genetic basis of craniofacial development and congenital malformations has been an area of investigation since the earliest understandings of heredity. Many genetic requirements were elucidated by targeted loss of function experiments for specific genes in multiple model organisms. Other genes have been identified by large scale mutagenesis forward genetic screens or even spontaneous mutations in animal colonies. An increasingly effective method is to directly apply next-generation sequencing efforts to affected humans and their biologic family members. The rapid advances in technology make sequencing ever more accessible, even to laboratory groups not traditionally considered to be human geneticists. The long-heralded ‘$1000 genome’ in which the entire human genome can be sequenced at sufficient depth for robust analysis is now readily available on a research basis.

The primary challenge then is not the identification of variant(s) which may contribute to a congenital craniofacial anomaly but, rather, assigning pathogenicity to a specific variant. One could attempt to do this by collecting multiple families with very similar phenotypic features and look for variants held in common across affected individuals from many families to identify variants that are not just shared by descent. This approach is, however, completely dependent on clinicians with phenotyping expertise to identify multiple families with key features, the difficulty of which is directly proportional to the rarity of the phenotype in the population. An alternative approach is to identify candidate variants within a family. While a trio approach with an affected proband and unaffected parents is most commonly used, phenotypes that seem to be segregating in a dominant fashion may require two trios as distantly related as possible within a family unless an affected family member with an apparent *de novo* variant is available.

A parallel approach is to generate a biological model of a suspected variant to truly demonstrate pathogenicity. This is tremendously facilitated by the recent advances in genome editing in both cellular and animal models. This approach will then also create an experimental substrate to understand the underlying molecular mechanism(s) and even potentially test therapeutic intervention strategies. A number of experimental options confront the researcher tasked with this challenge. Cellular models can be derived directly from the patients as fibroblasts or used in conjunction with cellular reprogramming techniques to create induced Pluripotent Stem Cells (iPSCs). iPSCs can then be used to create many different cell types. Isogenic iPSC lines can be made to rescue or recreate suspected pathogenic variants and test how the lines differ in experimental settings (Fatehullah, Tan, & Barker, 2016). Embryological models include frog, chick, and zebrafish. The relative merits of these have been extensively reviewed elsewhere, e.g., (Van Otterloo, Williams, & Artinger, 2016). Larger animal models can also be created including goats, pigs and monkeys with CRISPR-Cas9 editing (Cui et al., 2018; Hai, Teng, Guo, Li, & Zhou, 2014; Ni et al., 2014). We favor the mouse models for a combination of accessible genetics, relative cost, fast generation time, and physical resemblance to human biology as compared to many other genetic models. Here we illustrate both the power of this approach and some of the difficulties in modeling suspected human pathogenic variants in the mouse. We have used next-generation sequencing on a multi-generational family including two members with craniofacial anomalies. This analysis identified a compelling novel variant we suspected to be pathogenic. We then chose to model this variant and create a novel null allele of this gene in the mouse to test this prediction.

## RESULTS

### Whole exome sequencing of a family with a novel craniofacial syndrome

The proband was the 8 pound 13 ounce, 20 inch product of a 39 week gestation to a 39 year old Gravida 1, Para 0-1 woman and her 29 year old unrelated partner. Maternal gestational diabetes was noted at 8 weeks gestation but well controlled with insulin. The proband was born with a complete left sided cleft lip with cleft palate and right sided cleft of the primary palate. Prenatal fetal echocardiogram demonstrated a ventricular septal defect (VSD) in addition to the cleft lip with cleft palate. Postnatal echocardiogram showed a VSD with tetralogy of Fallot and a double orifice mitral valve. Neurologic examination was unremarkable. The early childhood clinical course was complicated by obstructive sleep apnea secondary to short mandible with glossoptosis. A large cisterna magna was incidentally identified during an airway MRI and has remained stable in subsequent imaging. At 14 years of age, advanced bone age (2.1 SD) was incidentally identified during hand films to evaluate the proband′s concerns about bent fingers.

The family history was significant as the proband′s father was also born with a left-sided unilateral complete cleft lip and cleft palate. He also had an atrial septal defect (ASD) repaired in early childhood and a reported mitral valve abnormality. Similar to the proband, the father was tall in stature and had large dysplastic ears. The proband has no biologic siblings and there is no other family history of cleft lip, cleft palate, or congenital heart defects.

At age 11 years, the proband, affected father, unaffected mother and unaffected paternal grandparents were enrolled in an institutionally funded investigational whole exome sequencing project for undiagnosed disorders (Fig. 1A). Alignment and variant detection were performed using the Broad Institute′s web-based Genome Analysis Toolkit (GATK; Genome Reference Consortium Build 37). The Golden Helix software suite was used for data filtering of VCF files containing a combined total of 107,135 variants. After quality control, 95,228 remaining variants were filtered for variants which were a) *de novo*, b) coding and non-synonymous and c) found only in the biologic father and transmitted to the proband. A single missense variant in *angiomotin like 1* (*AMOTL1*) 11:94532825, c.469C>T; p.Arg157Cys met all these criteria and presumably arose spontaneously in the father and was transmitted to the proband. The variant is in a region of the AMOTL1 protein which is highly conserved through many vertebrate species including chicken and frog (Fig. 1B). This variant is absent from the ExAC and gnomAD database at the time of writing; is considered ″deleterious″ by SIFT, ″disease causing″ by MutationTaster and ″probably damaging″ by PolyPhen-2; and the amino acid physiochemical difference is considered large with Grantham score of 180 (0 – 215). An independent analysis of the sequence was run with Ingenuity Variant Analysis (Qiagen), replicated the finding in the father and proband but also identified an *AMOTL1* missense variant c.712C>T, rs201051216 (MAF <0.001 in NHLBI ESP6500, European Americans) in the healthy mother that was transmitted to the son. Both *AMOTL1* variants were confirmed with Sanger sequencing in the proband and transmitting parents (Fig. 1C and data not shown). Further analysis focused on the unique paternal *de novo* c.469C>T variant since father and proband share key congenital anomalies.

**Figure 1.**
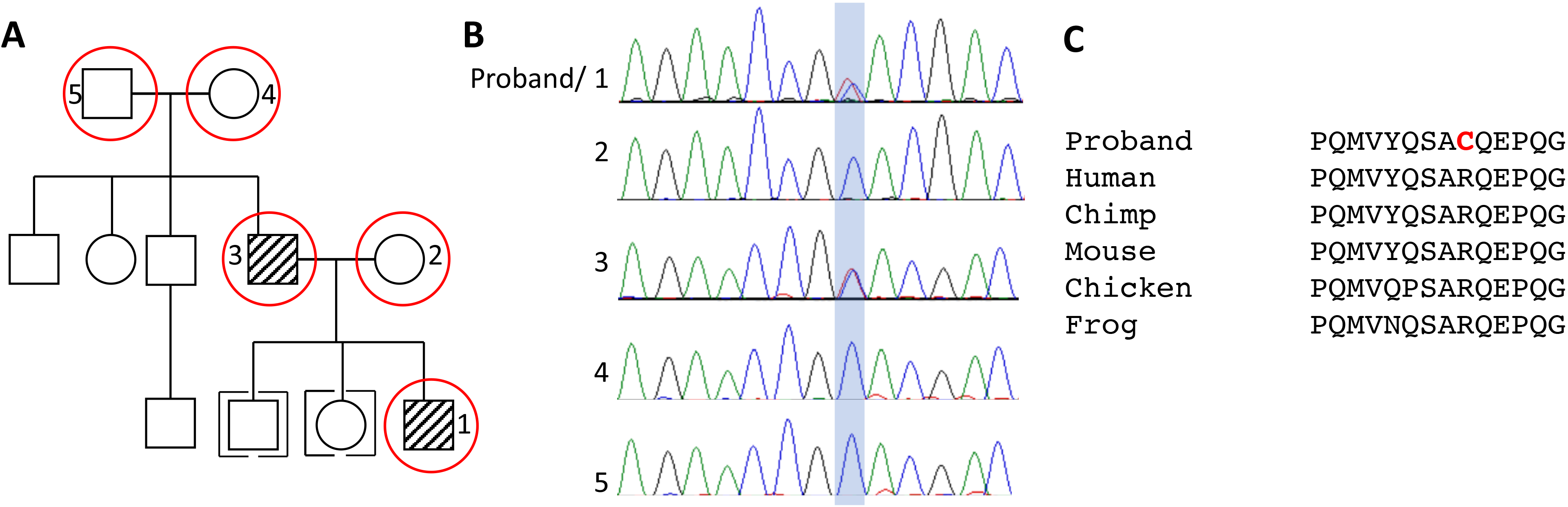
A novel congenital malformation syndrome. (A) Pedigree of proband (patient 1). Affected members are shown with the hash marks and the circled members were selected for exome sequencing. (B) The variant in *AMOTL1* affects a highly conserved arginine residue. (C) Sanger sequencing of indicated family members confirms the *AMOTL1* sequence variant is present in only patients 1 and 3.

### Novel mouse alleles of Amotl1

We took advantage of the ability to quickly make targeted mutations in the mouse with CRIPSR-Cas9 genome editing and constructed an allelic series of *Amotl1* with the goal of disrupting gene function and also recapitulating the R157C variant at the orthologous position in the mouse amino *add. Amotl1* had not been deleted in the mouse at the time of experimental design. We chose to keep and maintain two alleles from the CRISPR-Cas9 founder mice. *Amotl1*^*em1Rstot*^ (*Amotl1*^*R157C*^) is the arginine to cysteine coding change identified in the exome analysis. We also recovered a one base pair deletion *Amotlf^m2Rstot^ (Amotl1^01^)* which codes for a protein with 15 missense amino acids after position 155 and a stop codon (Fig. 2). We also recovered an eight base pair deletion *Amotl1*^*em3Rstot*^ (*Amotl1*^*D8*^) which codes for a protein with 34 missense amino acids after position 153 and a stop codon.

**Figure 2.**
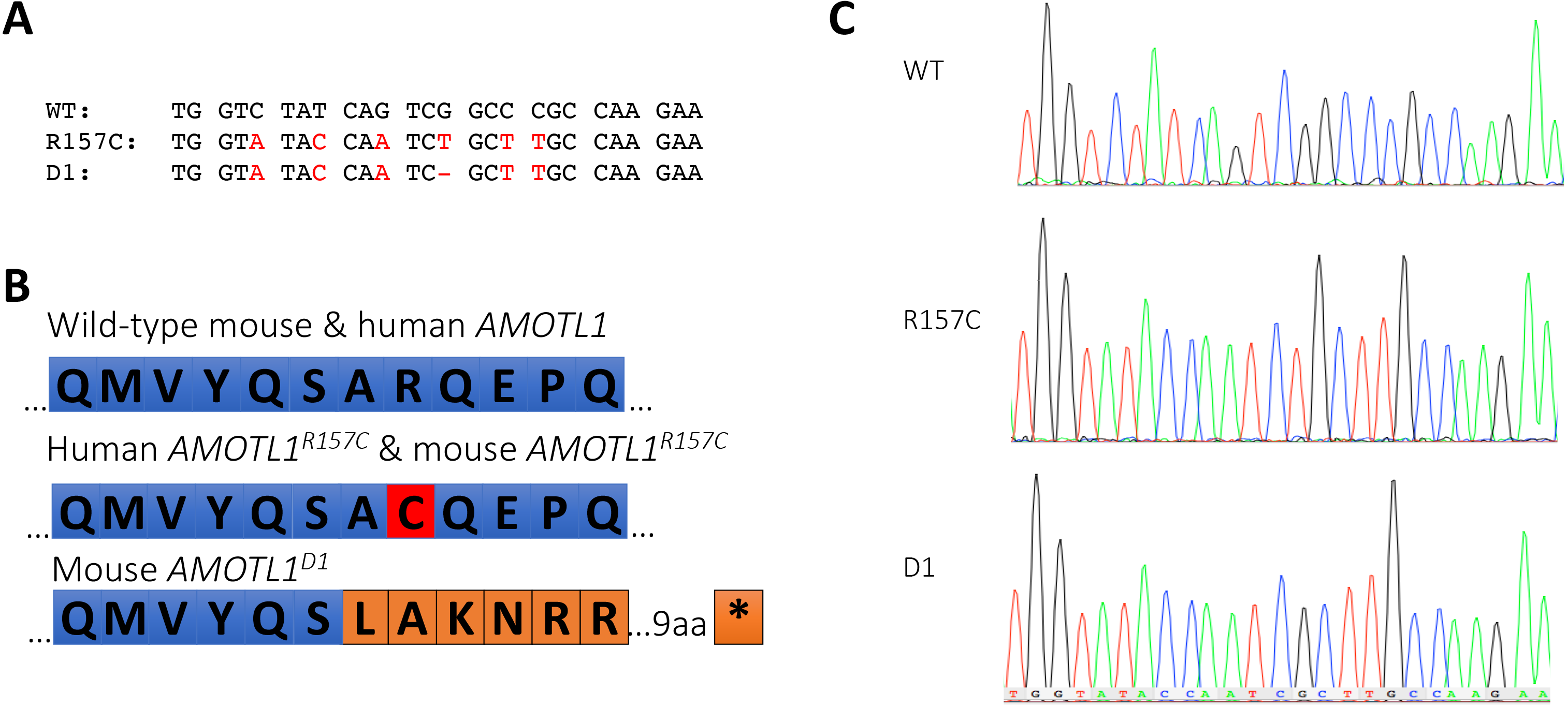
An allelic series of *Amotl1* in the mouse. (A) Wild-type nucleotide sequence of *Amotl1* around the desired single nucleotide C>T mutation (blue shaded box). Sequence analysis of *Amotl1*^*R157C*^ mice in this region indicates the desired genome edit was made along with other silent mutations (shown in red) to facilitate CRISPR guide stability and genotyping of modified mice. *Amotl1*^*D1*^ sequence is shown: (-) indicates the deleted nucleotide. (B) The predicted proteins for wild-type and each of the genome edits. The R157C variant is a single missense codon (red), while the D1 deletion changes codon usage to create 15 missense amino acids (orange) before a stop codon. (C) Sanger sequence validation of the sequence results schematized in (B).

We bred each of these alleles to test for viability (Table 1). We first noted that *Amotl1*^*R157C/wt*^ heterozygotes did not survive in Mendelian ratios at weaning as approximately one-third of the expected animals were not recovered. Surviving *Amotl1*^*R157C/wt*^ heterozygotes were intercrossed and we noted that the number of *Amotl1*^*R157C/R157C*^ homozygotes was substantially reduced at weaning and *Amotl1*^*R157C/wt*^ heterozygotes were again underrepresented from the remaining animals. We examined newborn litters at postnatal day (P)0 and P1 to begin to determine the stage of death and found the homozygotes were slightly under-represented at birth. An embryonic analysis from embryonic day (E)13.5-E18.5 showed no loss of *Amotl1*^*R157C/R157C*^ homozygotes or *Amotl1*^*R157C/wt*^ heterozygotes at organogenesis stages. We thus conclude that the *Amotl1*^*R157C*^ leads to incompletely penetrant lethality at perinatal stages. We next examined the survival of the *Amotl1*^*D1*^ deletion and found that animals homozygous for the *Amotl1*^*D1*^ deletion were viable and fertile as homozygotes with no significant deviation from Mendelian expectations. We found similar results with the*Amotl1*^*D8*^deletion and did not pursue this allele further.

**Table 1.** Survival of mice with targeted *Amotl1* alleles. * expectations for normal Mendelian ratios are shown parenthetically

|  | Wild-type | Heterozygous | Homozygous | Chi<sup>2</sup> |
| --- | --- | --- | --- | --- |
| *Amotl1* R157C x wt | 40 (31) | 22 (31) | N/A | 0.022 |
| *Amotl1* R157C intercross | 82 (50) | 107 (100) | 12 (50) | 1.69E-11 |
| *Amotl1* R157C (P0, P1) | 14 (6) | 7 (12) | 4 (6) | 0.056 |
| *Amotl1* R157C (E13.5-E18.5) | 29 (24) | 47 (48) | 22 (24) | 0.55 |
| *Amotl1* D1 intercross | 30 (25) | 47 (50) | 24 (25) | 0.54 |
| *Amotl1* D8 intercross | 26 (18) | 30 (36) | 15 (18) | 0.078 |

\*expectations for normal Mendelian ratios are shown parenthetically

The craniofacial and cardiac phenotypes in the human patients with the *Amotl1*^*R157C*^ variant both suggest mechanisms that might account for perinatal lethality in the mouse. We therefore examined the *Amotl1*^*R157C*^ mice for these phenotypes. We did not see any evidence of cleft palate in 24 *Amotl1*^*R157C/wt*^ embryos at E16.5-E18.5 and in 10 P0 *Amotl1*^*R157C/wt*^ pups. We also examined 9 *Amotl1*^*R157C/R157C*^ embryos at these stages and did not see signs of cleft palate in these animals either (Fig. 3A,B). Skeletal preparations, gross examinations and micro-CT imaging studies were performed but did not reveal any phenotypes to explain the perinatal lethality in either *Amotl1*^*R157C/wt*^ or *Amotl1*^*R157C/R157C*^ embryos (Fig. 3C-F). We also examined *Amotl1*^*R157C/wt*^ and *Amotl1*^*R157C/R157C*^ embryos for structural heart defects with a combination of whole mount and/or histological analysis with a special attention for septal and valve defects and outflow tract anomalies. Analysis of at least 16 *Amotl1*^*R157C/wt*^ and 6 *Amotl1*^*R157C/R157C*^ embryos at E16.5-E18.5 showed no evidence of outflow tract defects, septal defects or valvular anomalies. We also examined multiple *Amotl1*^*R157C/wt*^ mice at P28 and saw no evidence for heart phenotypes at this age either (Fig. 4).

**Figure 3.**
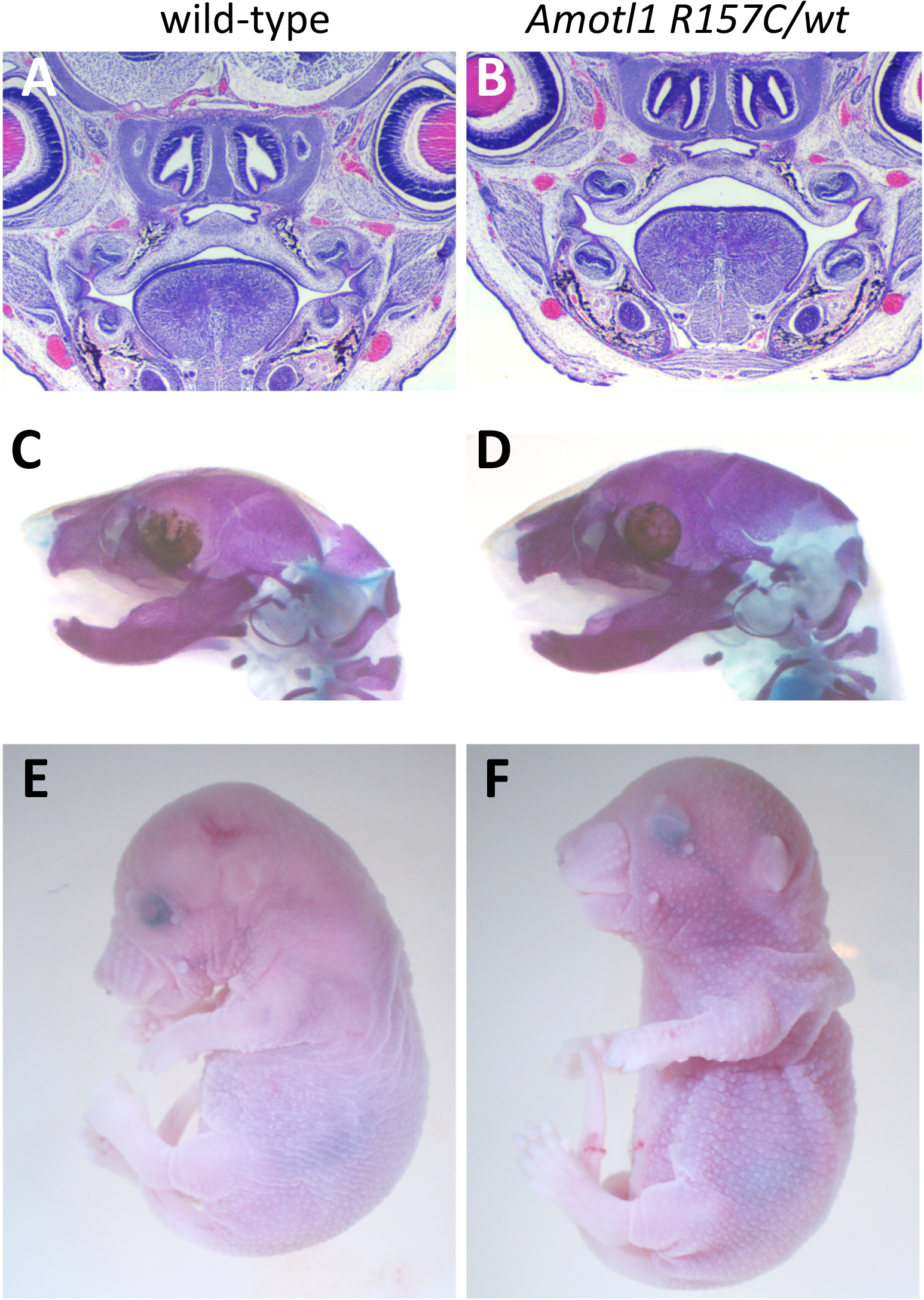
Mice carrying the *Amotl1*^*R157C*^ *variant are healthy and fertile*. Histological analysis (A,B), skeletal preparations (C,D) and whole mount analysis (E,F) do not reveal any phenotypes which allow us to explain the perinatal lethality in *Amotl1*^*R157C/wt*^ mice as compared to wild-type.

**Figure 4.**
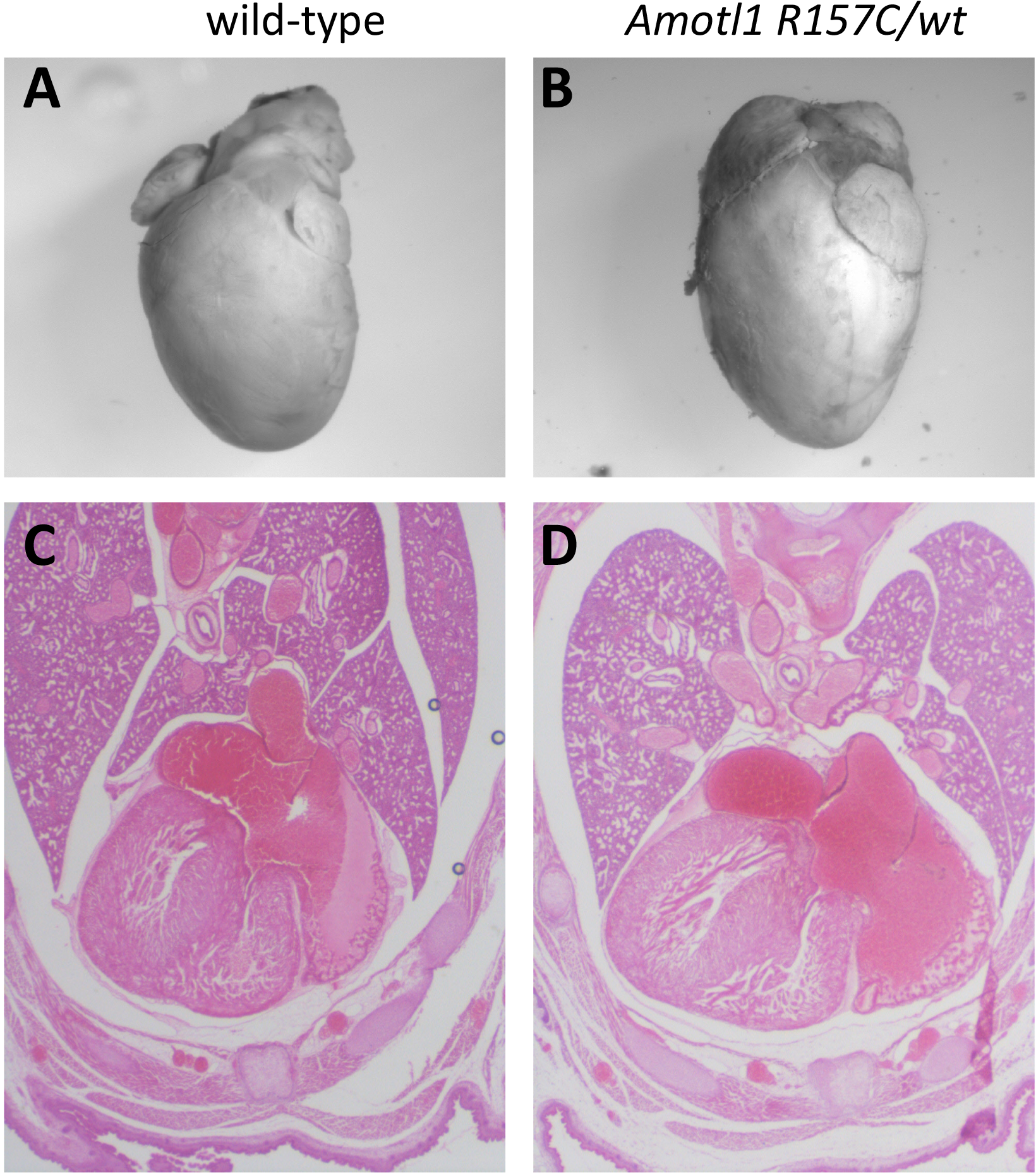
Cardiac development is normal in *Amotl1*^*R157C*^mice. Whole mount and histological analysis indicates cardiac development is relatively unaffected in *Amotl1*^*R157C/wt*^ mice.

The expression of a missense variant may lead to an unstable protein, so we over-expressed *in vivo* a myc-tagged AMOTL1 protein as well as a modified construct to recapitulate the *Amotl1*^*R157C*^ patient variant. Overexpression in HEK293T cells showed that the variant protein is produced by the cell but the Amotl1-R157C-myc variant runs as a larger protein species on the gel suggesting a post-translational modification may be present on the variant protein which is not found on control protein (Fig. 5A). We attempted to assay the levels of AMOTL1 protein in *Amotl1*^*D1/D1*^ mutant embryos. We were, however, unable to unambiguously detect AMOTL1 expression *in vivo* in a number of tissues where *Amotl1* expression is reported. Thus, we cannot definitively address the effects of these mutations on AMOTL1 levels *in vivo* but suspect the *Amotl1*^*D1*^ allele to be a loss of function or severely hypomorphic allele based on the amino acid sequence change. As the *Amotl1*^*D1*^ deletion creates a premature stop, we asked if the transcript for that variant was degraded by nonsense-mediated decay (He & Jacobson, 2015) and prepared cDNA from control and homozygous mutant embryos. We detected no change in the levels of RNA between mutant and wild-type (Fig. 5B).

**Figure 5.**
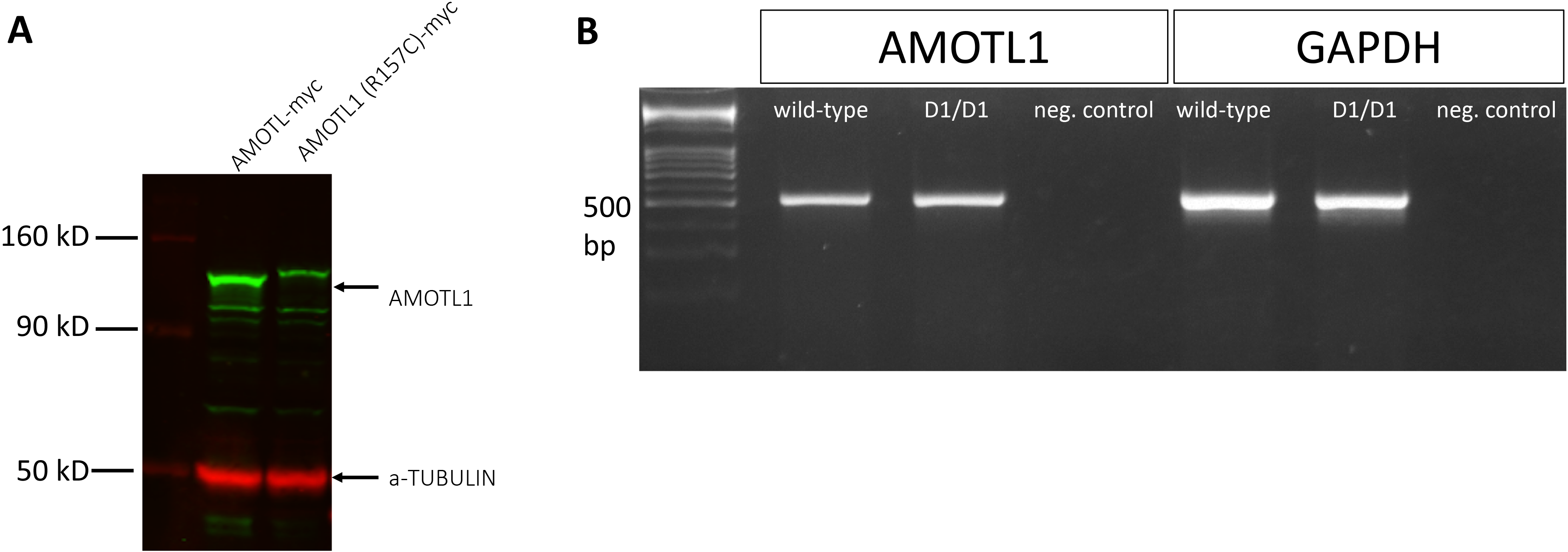
*Amotl1* variants do not affect significantly affect protein or mRNA levels. (A) Overexpression of AMOTL1 ^R157C^-myc indicates the coding change may allow a post-translational modification which changes the apparent size of the protein. Over a series of experiments, we do not notice a consistent and significant change in protein levels. (B) cDNA RT-PCR analysis shows that the premature stop codon in the Amotl1^D1^ allele does not trigger nonsense-mediated decay. *Gapdh* is shown as a control reaction.

## DISCUSSION

Here we illustrate one strategy to assess pathogenicity of variants identified in sequencing of a family with an inherited craniofacial anomaly. Exome sequencing of five members of the family suggested onec/e *novo* variant in *AMOTL1* may be the causal genetic lesion. Based on our preference for the mouse as the experimental model, we used CRISPR-Cas9 genome editing techniques to recreate the human variant and test the hypothesis this is the cause of the congenital anomaly. As a germline deletion of *Amotl1* has not been published to our knowledge, we took advantage of the possibility to also recover insertion/ deletion alleles from the transgenic founders. We conclude that *Amotl1* is not absolutely required in the mouse for survival to adulthood in a standard vivarium environment. We further show that the *Amotl1*^*R157C*^ missense variant identified in the human genetics studies significantly reduces viability of mice.

*Amotl1* is a member of the motin family of genes comprised of *angiomotin, angiomotin-like 1 (Amotl1)* and *angiomotin-like 2 (Amotl2)* (Bratt et al., 2002). Mice lacking *angiomotin* die around Ell.5 with vascular defects (Aase et al., 2007). Motin family members have been shown to interact with Wnt and Hippo signaling (Chan et al., 2011; Z. Li et al., 2012; Paramasivam, Sarkeshik, Yates, Fernandes, & McCollum, 2011; Ragni et al., 2017), but no germline deletion of *Amotl1* has been reported. Consistent with our conclusions, we do note that the international mouse phenotyping consortium (mousephenotype.org) has produced a gene trap allele of *Amotl1* and do not report any phenotypes in a thorough array of tests from 13 homozygous *Amotl1*^*tm1a(EUCOMM)Wtsi*^ mutants (Koscielny et al., 2014).

The interpretation of the data from the *Amotl1*^*R157C*^ allele is a bit more difficult. It is clear that the mutation compromises survival of the mouse and is probably acting as a dominant allele with incomplete penetrance. We were unable to determine the mechanism of death. The most obvious candidate systems based on time of death and the human proband phenotypes were craniofacial and cardiovascular development. However, we were not able to detect phenotypes in these tissues consistent with a perinatal lethality phenotype. We hypothesize that the *Amotl1*^*R157C*^ allele is a neomorphic allele which acts in a dominant negative fashion. We are not able then to definitively address the hypothesis that the *Amotl1*^*R157C*^ variant identified is related to the human pathology. We suspect that the human patients represent the more mild effects of a *AM0TL1*^*R157C*^ variant. Although we did not find comparable phenotypes between the human patients and mice, we do find it hard to believe we created the precisely same variant leading to a dominant effect on survival at random. However, as none of the phenotypes match between the human patients and mouse models, we are unable to make strong conclusions.

This study illustrates a few aspects of studies using animal models to assess human genetic findings. Next generation sequencing is now quite readily applicable to the study of human congenital malformations. This study was initiated a few years ago and utilized an exome sequencing approach focused on just the coding region of the genome. As sequencing technologies have evolved, unbiased whole genome sequencing is now becoming cost effective. Moreover, as the genome sequencing approach does not require the selective hybridization of the exonic sequences and avoids some of the biases from this library capture, the whole genome sequencing approach has been shown to generate more even coverage of the exome (Meienberg, Bruggmann, Oexle, & Matyas, 2016). We are now pursuing similar approaches to that described here in other families by directly employing whole genome sequencing at surprisingly affordable rates. We do, however, acknowledge that the genomics community is currently challenged to interpret the vast majority of non-coding variants that come out of a whole genome analysis. However, the data generated by this approach is a constant for each patient (with the caveat of acquired mosaic mutations), is easily stored, and can be periodically reanalyzed as our genomic understanding and capacity increases.

The amount of data in a whole genome sequencing experiment is immense but is relatively easy to manage in the context of rare congenital malformation studies. A crucial point of the experimental design is to include as many family members as practical and feasible in the analysis. The proband and parents (trio) are a normal minimum but any siblings (affected or unaffected) and extended family can be useful as well. This additional information allows for significant filtering and exclusion of identified variants. In the case of the family presented here, a dominant inheritance model was assumed. Thus, any variants not exclusive to the proband and his father can be eliminated in a first pass analysis. Another powerful tool is the ever-increasing number of genomes of control populations being deposited into public databases. This is especially true for the structural birth defects community. The genome aggregation database (gnomAD.broadinstitute.org) is one such resource and at the time of this writing has 123,136 exome sequences and 15,496 whole genome sequences (Lek et al., 2016). Other such efforts are also being made around the world. These projects allow us to begin to define the range of ″normal″ genetic variation in the adult human population. The individuals in the gnomAD project are known to not have severe pediatric disease. This method of collection means that in a first pass analysis of the results, any variant found with a minor allele frequency of greater than 1% or 0.1% in both the control databases and an affected individual can be initially classified as not causal. An even more aggressive strategy can potentially be applied where the presence of the allele in a control population to any degree suggests it may not be causal. Incomplete penetrance is of course a major caveat to this, but the first task in a genome analysis is to remove variants until a manageable number can be manually curated to formulate hypotheses. Datasets can always be re-analyzed with relaxed filters and constraints as necessary.

Recent complementary technologies make this a tremendously exciting time to be performing this kind of work with a wealth of tools at the disposal of the structural birth defect and craniofacial genetics research community. Animal models have been a valued experimental approach for a number of years but the advent of CRISPR mediated genome editing techniques means that conserved genes can be manipulated in model organisms to precisely model human variants. If the candidate gene in question has not previously been studied, the mechanics of CRISPR-Cas9 editing allow us to readily make an allelic series to create putative null alleles in the same experiment as the variant modeling. This is a fundamentally different approach than previous homologous recombination-based methodologies carried out in embryonic stem cells used to make chimeric founder animals. A potential challenge to this approach (especially relevant for craniofacial biology) is the modeling of a dominant negative allele. Modern medical care often allows humans with congenital craniofacial anomalies to thrive with surgical intervention. In the absence of such modalities, the laboratory mouse will die from many craniofacial anomalies preventing the researcher from establishing a stable transgenic line. An alternative approach is to perform the CRIPSR-Cas9 genome edits *in vitro* and, rather than let these animals be born to breed out the mutation, recover those treated embryos directly from the surrogate dam (an ‘FO CRISPR’ approach). A phenotypic and genomic analysis of these embryos can be performed to address hypotheses about genetic variants.

Conclusions about pathogenicity are much more convincing when the phenotypes of the animal model match the human model (DiStasio et al., 2017). These models then offer an experimental platform for detailed analysis of embryonic molecular mechanism as well as a tool to test any therapeutiv interventions that can be devised. Cases such as the one we report here represent examples where the interpretation is much more difficult and almost certainly can′t result in a clinically significant return of results to the family. The fact that the identical variant identified in exome analysis recapitulated in the mouse model results in unexplained lethality in mice suggests there clearly is some molecular pathogenesis. However, it is likely unwise to use this finding to direct clinical care. Rather, these represent tantalizing insights into structural birth defects biology that warrant further experimental effort.

## METHODS

### Patient sequencing and variant confirmation

Informed consent/assent was obtained from all subjects according to Cincinnati Children′s Hospital Medical Center (CCHMC) institutional review board protocol #2012-0203. All methods and experimental protocols were carried out in accordance with relevant guidelines and regulations and with approval from the CCHMC Institutional Biosafety Committee. Following consent, whole blood was collected. Library generation, exome enrichment, sequencing, alignment and variant detection were performed in the CCHMC Genetic Variation and Gene Discovery Core Facility (Cincinnati, OH). Briefly, sheared genomic DNA was enriched with NimbleGen EZ Exome V2 kit (Madison, WI). The exome library was sequenced using lllumina′s Hi Seq 2000 (San Diego, CA). Alignment and variant detection was performed using the Broad Institute′s web-based Genome Analysis Tookit (GATK)(McKenna et al., 2010). All analyses were performed using Genome Reference Consortium Build 37.

### Variant Filtering and Pathogenicity Assessment

Quality control and data filtering were performed on VCF files in Golden Helix′s SNP and Variation Suite (Bozeman, MT) as well as Ingenuity Variant Analysis (Qiagen, Germany). Non-synonymous coding variants were compared to three control databases, including NHLBI′s ESP6500 exome data (Fu et al., 2013), the 1000 genomes project (Genomes Project et al., 2010), EXAC Browser (Karczewski et al., 2017) and an internal CCHMC control cohort (Patel et al., 2014). Remaining variants were subject to autosomal recessive analysis with emphasis on homozygous recessive variants found in the region of homozygosity identified by SNP microarray. The identified variant was compared to known disease genes in the OMIM and Human Gene Mutation (HGMD) (Stenson et al., 2014) databases, and to reported variants in dbSNP (Sherry et al., 2001) and the Exome Variant Server. The variant was also analyzed using Interactive Biosoftware′s Alamut v2.2 (San Diego, CA) to determine location of mutation within a protein domain, the conservation of the amino acid, the Grantham score (Grantham, 1974) and the designation of the mutation by three existing in-silico software tools, SIFT (Li, Gui, Kwan, Bao, & Sham, 2012), Polyphen (Adzhubei et al., 2010) and Mutation Taster (Schwarz, Rodelsperger, Schuelke, & Seelow, 2010).

### Mouse allele generation

CRISPR guides for *Amotl1* were evaluated using the Fusi (Benchling.com) and Moreno-Mateos (Crisprscan.org) algorithms. Potential guide RNA (gRNA) sequences were selected and ordered as complementary oligonucleotide pairs with BbsI over-hangs (IDT, Coralville, IA). These were ligated into the pSpCas9(BB)-2A-GFP (px458) vector and transfected into MK4 cells at low confluence using the Lipofectamine2000 transfection reagent (Thermo Fisher Scientific, Massachusetts). Cells were harvested 48 h after transfection and genomic DNA was isolated and used with the Surveyor mutation detection kit (IDT) in order to test gRNA cutting efficiency. As a control, cutting efficiencies of potential guides were compared with that of a previously-published mTet2 gRNA. Cas9 and gRNAs were injected into C57BL/6N zygotes (Taconic) by the CCHMC Transgenic Core. Potential founders were validated with Sanger sequencing of tail DNA and subsequently maintained on a C57BL/6J (Jackson Labs) background. pSpCas9(BB)-2 A-GFP (PX458) was a gift from Feng Zhang (Addgene plasmid # 48138). All alleles are available to the research community upon publication.

### Animal husbandry

All animals were housed under an approved protocol in standard conditions. All euthanasia and subsequent embryo or organ harvests were preceded by Isoflurane sedation. Euthanasia was accomplished via dislocation of the cervical vertebrae. For embryo collections, noon of the day of vaginal plug detection was designated as E0.5.

### Histological analysis

Tissue samples were fixed in Bouin′s fixative solution. Samples were then paraffin embedded, sectioned at 6μm for adult tissue and 10μm for embryonic tissue, and processed through hematoxylin and eosin (H&E) or Nissl staining. Sections were sealed using Cytoseal Mounting Medium (ThermoFisher-Scientific). All paired images are shown at the same magnification.

### Amotl1 overexpression and cDNA analysis

An *Amotl1* expression construction encoding an AMOTL1-myc fusion protein tag was purchased from Origene (#RC207693, Rockville, MD). Site directed mutagenesis to recreate the R157C variants was performed with QuikChange Lightning Site-Directed Mutagenesis kit (Agilent, Santa Clara, CA). HEK293T cells were transfected with standard protocols. Western immunoblotting was performed with standard protocols and a primary polyclonal rabbit antibody (PA5-42267 from ThermoFisher). RNA was extracted and purified from *Amotl1 ^D1^* animals and wild-type controls with TRIZOL reagent. cDNA was prepared with Superscript III reverse transcriptase and random hexamer primers.

**Table 2.**
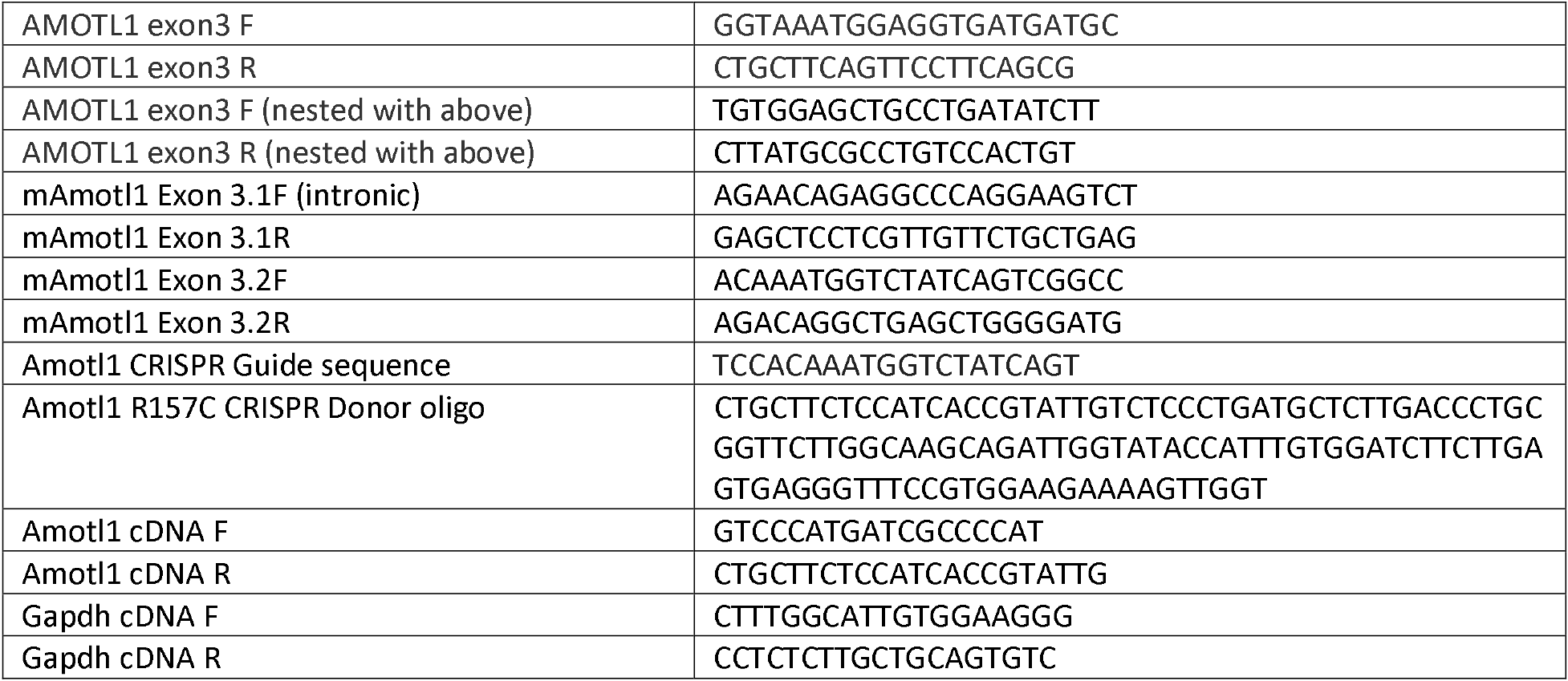
**PCR Primers.**

## ACKNOWLEDGEMENTS

We are grateful to Yueh-Chiang Hu and the CCHMC Transgenic Animal and Genome Editing Core for advice and technical assistance, and to Katherine Yutzey for extensive discussion of the cardiovascular analysis prior to publication. Funding for this study was provided by the Cincinnati Children’s Research Foundation and the CCHMC Center for Pediatric Genomics.

## REFERENCES

Aase, K., Ernkvist, M., Ebarasi, L., Jakobsson, L., Majumdar, A., Yi, C., … Holmgren, L. (2007). Angiomotin regulates endothelial cell migration during embryonic angiogenesis. Genes Dev, 21(16), 2055–2068. doi:10.1101/gad.432007

Adzhubei, I. A., Schmidt, S., Peshkin, L., Ramensky, V. E., Gerasimova, A., Bork, P.,… Sunyaev, S. R. (2010). A method and server for predicting damaging missense mutations. Nat Methods, 7(4), 248–249. doi:10.1038/nmeth0410-248

Bratt, A., Wilson, W. J., Troyanovsky, B., Aase, K., Kessler, R., Van Meir, E. G., & Holmgren, L. (2002). Angiomotin belongs to a novel protein family with conserved coiled-coil and PDZ binding domains. Gene, 298(1), 69–77.

Chan, S. W., Lim, C. J., Chong, Y. F., Pobbati, A. V., Huang, C., & Hong, W. (2011). Hippo pathway-independent restriction of TAZ and YAP by angiomotin. J Biol Chem, 286(9), 7018–7026. doi:10.1074/jbc.C110.212621

Cui, Y., Niu, Y., Zhou, J., Chen, Y., Cheng, Y., Li, S.,… Ji, W. (2018). Generation of a precise Oct4-hrGFP knockin cynomolgus monkey model via CRISPR/Cas9-assisted homologous recombination. Cell Res, 28(3), 383–386. doi:10.1038/cr.2018.10

DiStasio, A., Driver, A., Sund, K., Donlin, M., Muraleedharan, R. M., Pooya, S.,… Stottmann, R. W. (2017). Copb2 is essential for embryogenesis and hypomorphic mutations cause human microcephaly. Hum Mol Genet, 26(24), 4836–4848. doi:10.1093/hmg/ddx362

Fatehullah, A., Tan, S. H., & Barker, N. (2016). Organoids as an in vitro model of human development and disease. Nat Cell Biol, 18(3), 246–254. doi:10.1038/ncb3312

Fu, W., O’Connor, T. D., Jun, G., Kang, H. M., Abecasis, G., Leal, S. M.,… Akey, J. M. (2013). Analysis of 6,515 exomes reveals the recent origin of most human protein-coding variants. Nature, 493(7431), 216–220. doi:10.1038/naturell690

Genomes Project, C., Abecasis, G. R., Altshuler, D., Auton, A., Brooks, L. D., Durbin, R. M.,… McVean, G. A. (2010). A map of human genome variation from population-scale sequencing. Nature, 467(1319), 1061–1073. doi:10.1038/nature09534

Global strategies to reduce the health care burden of craniofacial anomalies: report of WHO meetings on international collaborative research on craniofacial anomalies. (2004). Cleft Palate CraniofacJ, 41(3), 238–243. doi:10.1597/03-214.1

Grantham, R. (1974). Amino acid difference formula to help explain protein evolution. Science, 185(4154), 862–864.

Hai, T., Teng, F., Guo, R., Li, W., & Zhou, Q. (2014). One-step generation of knockout pigs by zygote injection of CRISPR/Cas system. Cell Res, 24(3), 372–375. doi:10.1038/cr.2014.11

He, F., & Jacobson, A. (2015). Nonsense-Mediated mRNA Decay: Degradation of Defective Transcripts Is Only Part of the Story. Annu Rev Genet, 49, 339–366. doi:10.1146/annurev-genet-112414-054639

Karczewski, K. J., Weisburd, B., Thomas, B., Solomonson, M., Ruderfer, D. M., Kavanagh, D.,… MacArthur, D. G. (2017). The ExAC browser: displaying reference data information from over 60 000 exomes. Nucleic Acids Res, 45(D1), D840–D845. doi:10.1093/nar/gkw971

Koscielny, G., Yaikhom, G., Iyer, V., Meehan, T. F., Morgan, H., Atienza-Herrero, J.,… Parkinson, H. (2014). The International Mouse Phenotyping Consortium Web Portal, a unified point of access for knockout mice and related phenotyping data. Nucleic Acids Res, 42(Database issue), D802–809. doi:10.1093/nar/gkt977

Lek, M., Karczewski, K. J., Minikel, E. V., Samocha, K. E., Banks, E., Fennell, T.,… Exome Aggregation, C. (2016). Analysis of protein-coding genetic variation in 60,706 humans. Nature, 536(7616), 285–291. doi:10.1038/naturel9057

Li, M. X., Gui, H. S., Kwan, J. S., Bao, S. Y., & Sham, P. C. (2012). A comprehensive framework for prioritizing variants in exome sequencing studies of Mendelian diseases. Nucleic Acids Res, 40(7), e53. doi:10.1093/nar/gkrl257

Li, Z., Wang, Y., Zhang, M., Xu, P., Huang, H., Wu, D., & Meng, A. (2012). The Amotl2 gene inhibits Wnt/beta-catenin signaling and regulates embryonic development in zebrafish. J BiolChem, 287(16), 13005–13015. doi:10.1074/jbc.M112.347419

McKenna, A., Hanna, M., Banks, E., Sivachenko, A., Cibulskis, K., Kernytsky, A.,… DePristo, M.A. (2010). The Genome Analysis Toolkit: a MapReduce framework for analyzing next-generation DNA sequencing data. Genome Res, 20(9), 1297–1303. doi:10.1101/gr.107524.110

Meienberg, J., Bruggmann, R., Oexle, K., & Matyas, G. (2016). Clinical sequencing: is WGS the better WES? Hum Genet, 135(3), 359–362. doi:10.1007/s00439-015-1631-9

Ni, W., Qiao, J., Hu, S., Zhao, X., Regouski, M., Yang, M.,… Chen, C. (2014). Efficient gene knockout in goats using CRISPR/Cas9 system. PLoS One, 9(9), e106718. doi:10.1371/journal.pone.0106718

Paramasivam, M., Sarkeshik, A., Yates, J. R. 3rd, Fernandes, M. J., & McCollum, D. (2011). Angiomotin family proteins are novel activators of the LATS2 kinase tumor suppressor. Mol Biol Cell, 22(19), 3725–3733. doi:10.1091/mbc.Ell-04-0300

Patel, Z. H., Kottyan, L. C., Lazaro, S., Williams, M. S., Ledbetter, D. H., Tromp, H.,… Kaufman, K. M. (2014). The struggle to find reliable results in exome sequencing data: filtering out Mendelian errors. Front Genet, 5,16. doi:10.3389/fgene.2014.00016

Ragni, C. V., Diguet, N., Le Garrec, J. F., Novotova, M., Resende, T. P., Pop, S.,… Meilhac, S. M. (2017). Amotl1 mediates sequestration of the Hippo effector Yap1 downstream of Fat4 to restrict heart growth. Nat Commun, 8,14582. doi:10.1038/ncommsl4582

Schwarz, J. M., Rodelsperger, C., Schuelke, M., & Seelow, D. (2010). MutationTaster evaluates disease-causing potential of sequence alterations. Nat Methods, 7(8), 575–576. doi:10.1038/nmeth0810-575

Sherry, S. T., Ward, M. H., Kholodov, M., Baker, J., Phan, L., Smigielski, E. M., & Sirotkin, K. (2001). dbSNP: the NCBI database of genetic variation. Nucleic Acids Res, 29(1), 308–311.

Stenson, P. D., Mort, M., Ball, E. V., Shaw, K., Phillips, A., & Cooper, D. N. (2014). The Human Gene Mutation Database: building a comprehensive mutation repository for clinical and molecular genetics, diagnostic testing and personalized genomic medicine. Hum Genet, 133(1), 1–9. doi:10.1007/s00439-013-1358-4

Van Otterloo, E., Williams, T., & Artinger, K. B. (2016). The old and new face of craniofacial research: How animal models inform human craniofacial genetic and clinical data. Dev Biol, 415(2), 171–187. doi:10.1016/j.ydbio.2016.01.017

